# cycle_finder: *de novo* analysis of tandem and interspersed repeats based on cycle-finding

**DOI:** 10.1101/2023.07.17.549334

**Authors:** Yoshiki Tanaka, Rei Kajitani, Takehiko Itoh

## Abstract

Repeat sequences in the genome can be classified into interspersed and tandem repeats, both of which are important for understanding genome evolution and important traits such as disease. They are also noteworthy as regions of high frequency of genome rearrangement in somatic cells and high inter-individual diversity. Existing repeat detection tools have limitations in that they targets only one of the two types and/or require reference sequences. In this study, we developed a novel tool: cycle_finder, which constructs a graph structure (de Bruijn graph) from low-cost short-read data and constructs units of both types of repeats. The tool can detect cycles with branching and corresponding tandem repeats, and can also construct interspersed repeats by exploring non-cycle subgraphs. Furthermore, it can estimate sequences with large copy-number differences by using two samples as input. Benchmarking with simulations and actual data from the human genome showed that this tool had superior recall and precision values compared to existing methods. In a test on the roundworm data, in which large-scale deletions occur in somatic cells, the tool succeeded in detecting deletion sequences reported in previous studies. This tool is expected to enable low-cost analysis of repeat sequences that were previously difficult to construct.

## Introduction

Repeat sequences often account for a large proportion of eukaryotic genomes and are classified into interspersed repeats and tandem repeats. Interspersed repeats, represented by transposons, have been suggested to have a major impact on genome evolution (Senft and Macfarlan, 2021) and many cases related to important traits such as diseases have been reported (Payer and Burns, 2019). Tandem repeats are represented by satellite sequences, and this centromere-related sequence plays a function in chromosome segregation during cell division and is also associated with disease (Black and Giunta, 2018; Barra and Fachinetti, 2018). It has been suggested that human centromere regions have high divergence between individuals (Suzuki *et al*., 2020) and are important for understanding genomic diversity. Tandem repeats are also associated with somatic mutations, with reports of somatic deletion of tandem repeat-rich regions in various lineages (Wang and Davis, 2014). A representative cases are those of hagfishes (Kubota *et al*., 2001) and roundworms(Wang *et al*., 2012). The study of important topics such as evolution, diversity, disease, and somatic mutation requires a comprehensive analysis for both interspersed and tandem repeats.

In general, repeat sequence regions are difficult to construct by *de novo* assembly. In humans, the complete genome sequence has been determined (Nurk *et al*., 2022) A region rich in tandem repeat sequences of approximately 200 Mbp was newly determined. In this case, multiple types of long-read data and a proprietary pipeline were applied, indicating that there are many regions that cannot be constructed by short-read or Sanger sequencing. When analyzing for somatic mutations and inter-individual diversity, it is desirable to target multiple samples, but it is costly to prepare ample long-read data sets for all samples. The cost of sequencing short reads is small (Jeon *et al*., 2019)), and if repeat sequences can be efficiently constructed from short reads, it is expected to facilitate research on those topics.

A specialized tool for the construction of tandem repeats is DExTAR (Fertin *et al*., 2014) and MIXTAR (Fertin *et al*., 2015), TAREAN (Novák *et al*., 2017) etc., and those targeting interspersed repeats include RepARK (Koch *et al*., 2014), Repdenovo (Chu *et al*., 2016) and others. No tool exists to our knowledge that comprehensively constructs both repeat sequences; MIXTAR is an improved version of DExTAR, but has individual limitations, such as the need for long reads and an upper limit of 100 bp for detectable unit length. Tools for detecting repeat sequences on reference sequences are also available, such as the Tandem Repeat Finder (TRF) (Benson, 1999) However, these are not suitable for analyzing repeat sequences that fail to be constructed at the time of draft genome assembly. There is also a lack of tools to perform analyses while comparing multiple data sets (e.g., germline and somatic cells). An example of a possible analysis would be the extraction of large differences in frequency between two samples.

In this study, we developed a tool: cycle_finder, which allows *de novo* construction of both tandem repeats and interspersed repeats, and copy number comparison between two samples. It constructs a data structure called a de Bruijn graph from low-cost short-read data, and first constructs tandem repeat units by detecting cycles in the graph. The tool then constructs a subgraph excluding the cycles and constructs interspersed repeat sequences within the subgraph, and when two samples are input, the tool extracts and analyzes sequences with particularly large differences in the number of occurrences between samples. This tool enables efficient and exhaustive analysis of repeat sequences and is expected to promote research on somatic mutation and diversity.

## Methods

### Overview of cycle_finder

The flow of this method can be broadly categorized as follows (Figure 1): (1) extraction of high frequency *k*-mer, (2) repeat detection from de Bruijn graph, (3) clustering, and (4) copy number estimation. Although this method was developed to perform comparative genome analysis of two types of genomes, it can be applied to a single type of genome. In the following, we refer to the method that compares two types of data and detects repeat sequences as “compare mode” and the method that detects high frequency repeat sequences from only one type of data as “single mode.”

**Figure 1.**
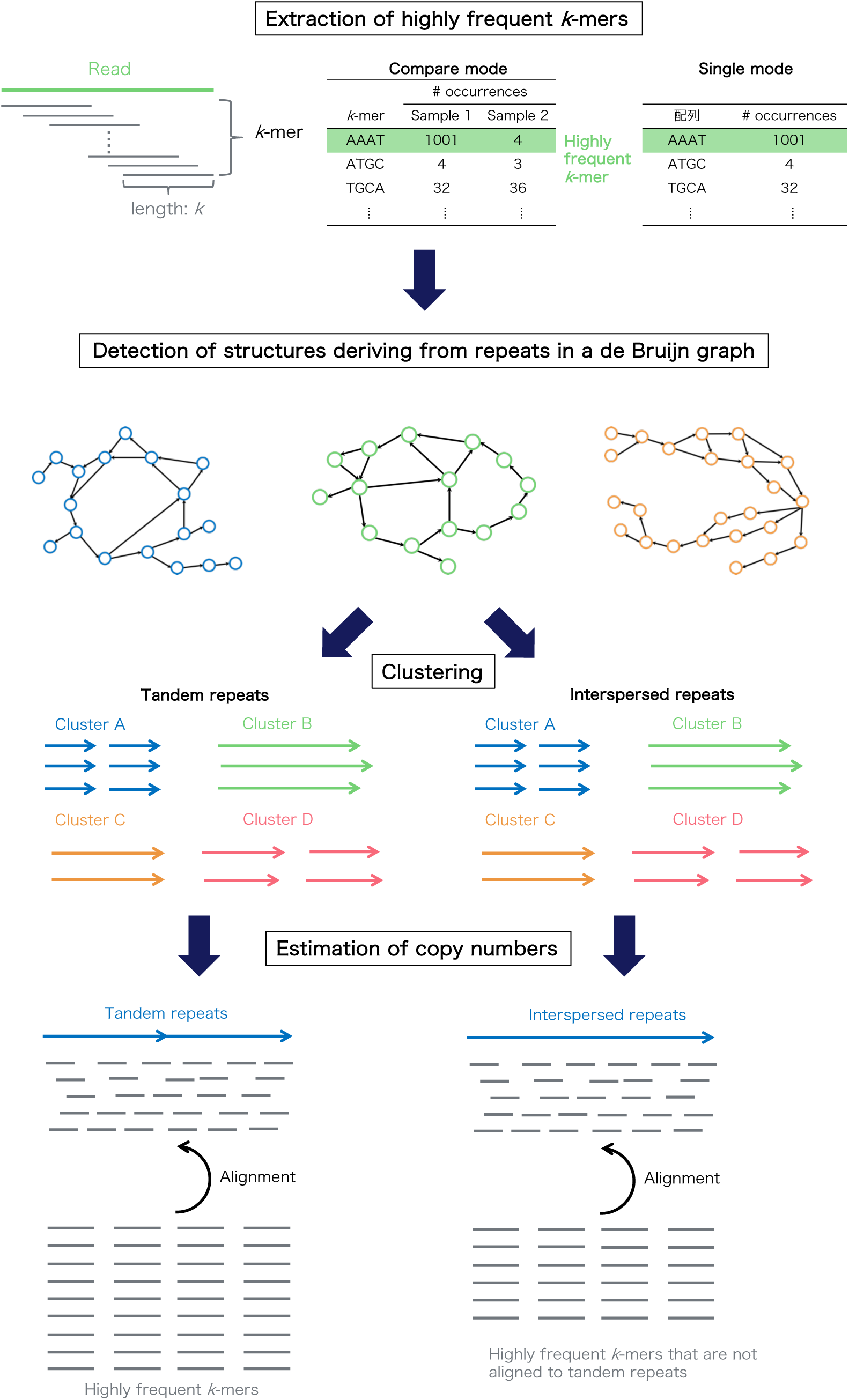
Overview of cycle_finder.

### External tools to be utilized

The method uses existing tools in alignment and clustering, which are listed below.

(1) BLASTN: a tool for aligning similar sequences (Camacho *et al*., 2009)
(2) CD-HIT: a tool for clustering similar sequences (Li *et al*., 2012)
(3) Jellyfish: a tool for extracting *k*-mer from short reads and counting the number of *k*-mer occurrences (Marçais and Kingsford, 2011)
(4) Tandem Repeat Finder: a tool to detect tandem repeats in a sequence. (Benson, 1999)

### Extraction of high-frequency *k*-mer

In order to extract high frequency *k*-mer derived from repeat sequences, *k*-mer with more than *α* occurrences in the genome are extracted in compare mode, and *k*-mer with more than α occurrences in the genome are extracted in single mode. The smaller the value of α, the more sensitive the detection, but the smaller the value of α, the longer the computation time increases. The default value is set to *α* = 10. In addition, Jellyfish, an existing tool, is used to obtain *k*-mer from short reads. The canonical option is used to treat complementary strand sequences as the same sequence.

### Tandem repeat detection

Detect tandem repeats from extracted high-frequency *k*-mer.

#### (1) Construction of de Bruijn graphs

Here, we consider the construction of de Bruijn graphs from the extracted set of *k*-mers. de Bruijn graphs are graphs that consider a k-mer as a node and show the edges connecting two *k*-mers when they have (*k*−1)-characters in common sequence (Figure 2). In this case, the graph is a directed graph consisting only of nodes (vertices) and edges with an orientation that connects the nodes. However, the graph treated in this method differs from de Bruijn graphs in the strict sense of the term, and has the following properties.

i. A *k*-mer and its complementary strand are treated identically.
ii. The graph is a directed graph and represents an edge from k-mer1 to *k*-mer2 when the back (*k*−1) character of *k*-mer1 and the front (*k*−1) character of *k*-mer2 have (*k*−1) characters in common.
iii. Each node has a value of *k*-mer occurrences.

**Figure 2.**
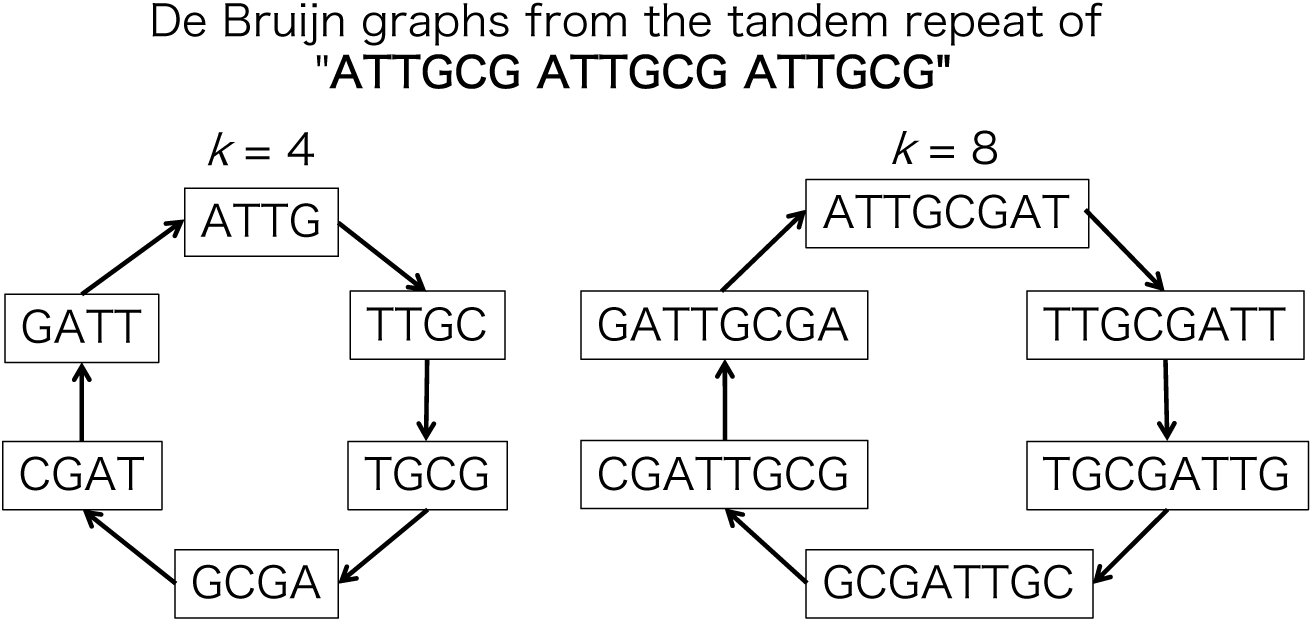
Example of de Bruijn graph in tandem repeat.

#### (2) Properties of tandem repeats in de Bruijn graphs

In de Bruijn graphs, tandem repeats exhibit the property of forming cycles (Figure 2). This property does not depend on the value of *k* in relation to the unit length of the tandem repeats. This property is used to detect tandem repeats by searching for cycles in de Bruijn graphs.

#### (3) Search for cycles

The algorithm for cycle search was implemented based on Johnson’s algorithm (Johnson, 1975). It is an algorithm for cycle search used in DExTAR (Fertin et al., 2014) and MIXTAR (Fertin et al., 2015), which uses depth-first search to search for cycles. Depth-first search is Figure 3 When considering a path in a graph, as shown in Figure 3, the search method selects one of the branches and searches deeper nodes until it cannot search any more, and when it cannot search any more, it returns to the other branch that is closest to it. When the search becomes impossible, it returns to the other side of the nearest branch. In this method, the *k*-mer (node) with the highest number of occurrences in the branching is preferentially searched.

**Figure 3.**
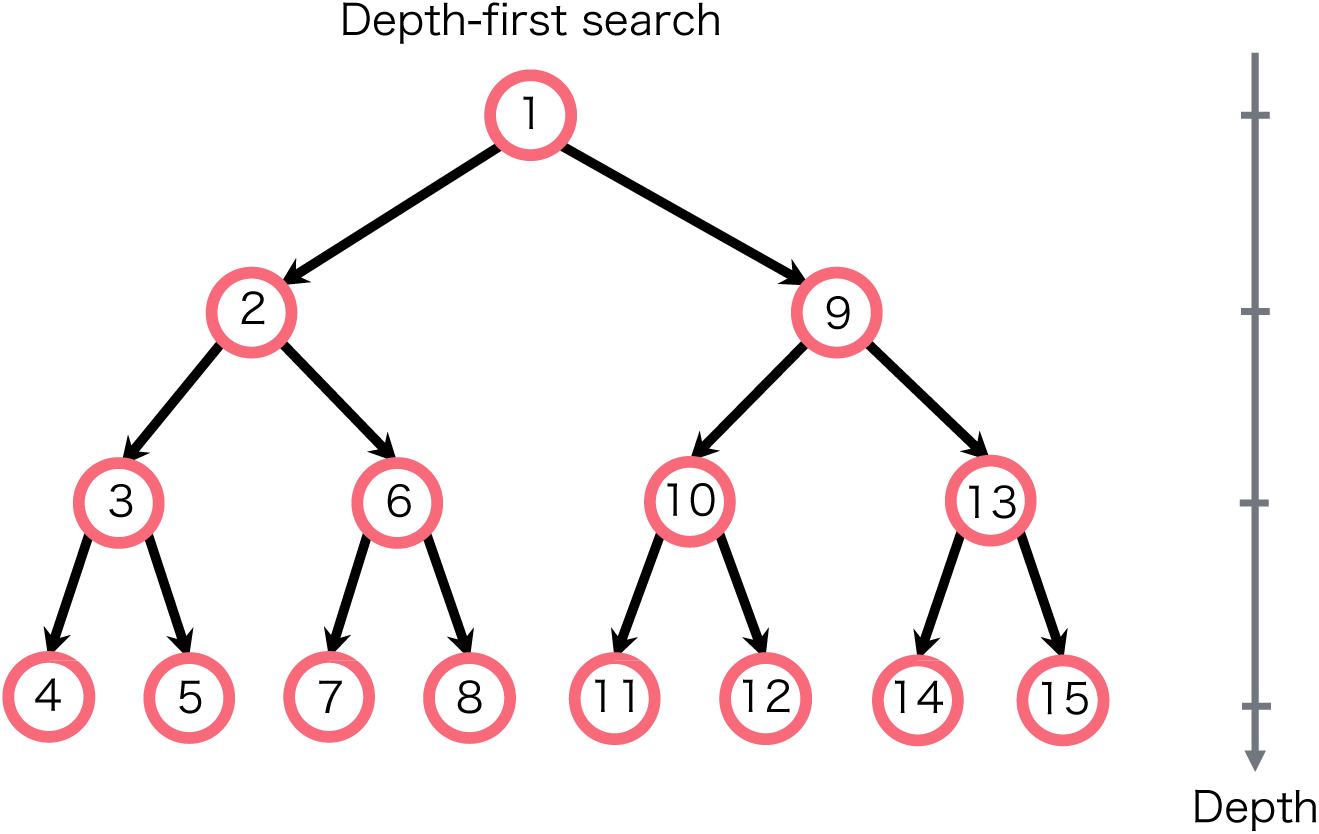
Search order for depth-first search.

If the number of nodes is *n*, the number of edges is *e*, and the number of cycles is *c*, the time cost of cycle search is expressed in Equation 1 (Johnson, 1975).

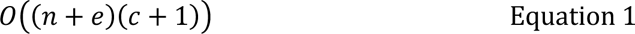

In the previous step, high-frequency *k*-mer extraction was performed. If the threshold value *α* set at this time is small, *n, e,* and *c* in the above equation increase, resulting in a significant increase in computation time. In addition, since we are considering *k-*mer in the genome, the number of these is very large, so the value of *α* must be raised to find all the cycles, which greatly reduces the detection sensitivity. To address this problem, we set the following parameters to search for cycles heuristically while keeping *α* small.

*l*: Upper limit of tandem repeat length to be detected [default value: 1,000]

*n*: Upper limit on the number of nodes to search [default value: 10,000].

*d*: Upper limit of branching depth to search [default value: 10]. The following is a description of the cycle search procedure.

i. Extract *k*-mer where the product of the number of incoming edges and the number of outgoing edges is greater than 1. The cycle in the de Bruijn graph is Figure 4 The cycles in the de Bruijn graph can be classified into two categories: those with branches and those without branches, as shown in Figure 4. In the former case, the search is limited to *k*-mers with branches, which requires less computation time, thus *k*-mers with branches are extracted.
ii. Depth-first search is performed starting from the *k*-mer with the branching point. The order of search at branching points is determined by the number of occurrences of the next *k*-mer, with priority given to those with the highest number of occurrences. If it returns to the starting point, it is stored as a cycle and the search continues. However, if the path returns to a node other than the starting point, it returns to the previous branch and continues the search without storing the path. If all paths are explored or if any of the set parameters *l, n, or d* are exceeded, the search is terminated and the starting node is removed from the graph.
iii. Perform a search step for ii starting at the next *k*-mer.
iv. If the search ends with a branching *k-*mer as the starting point, a depth-first search is performed starting at the *k-*mer where the number of incoming edges and the number of outgoing edges are both 1 in order to find a cycle for the branch, and the path is stored as a cycle if it returns to the starting point.

**Figure 4.**
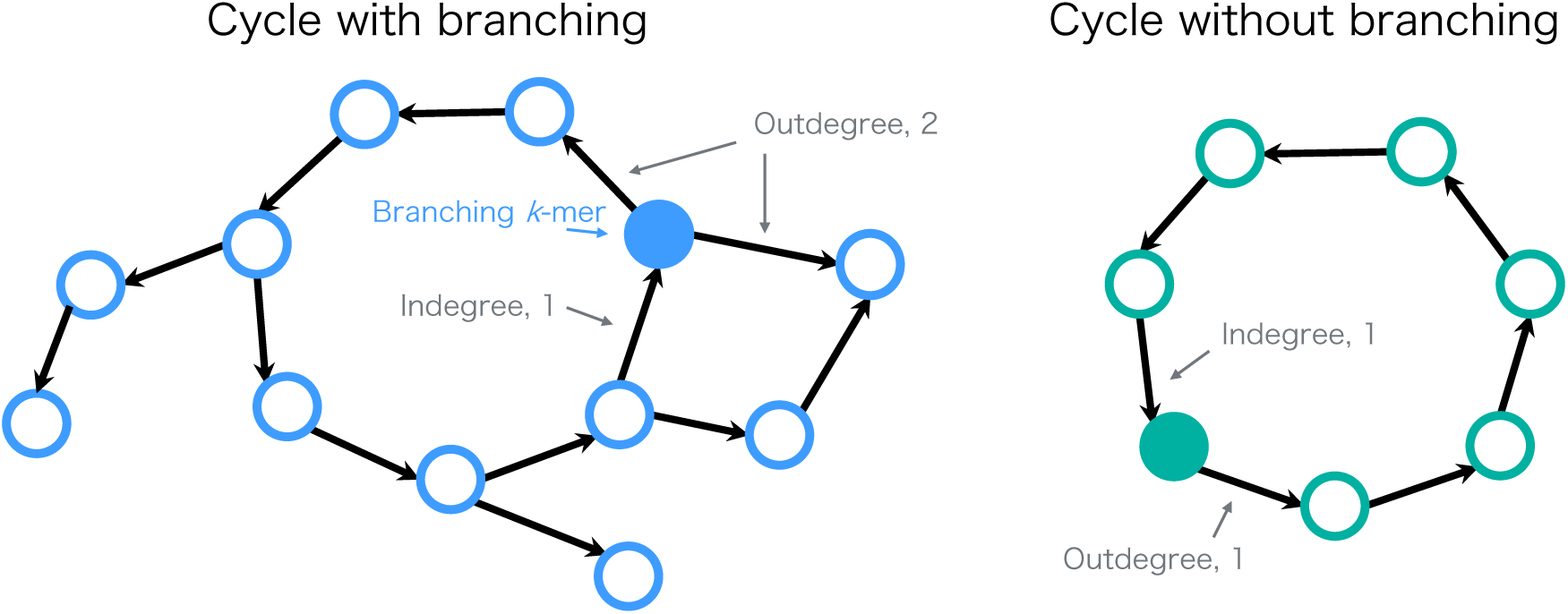
Two types of cycles.

#### (4) Removal of repeats containing multiple units by TRF

Some of the tandem repeats detected in the cycle may contain multiple tandem repeat sequences in a single unit that differ by a few bases due to mutations. This is especially common in microsatellites with short unit lengths. Tandem Repeat Finder (TRF) (Benson, 1999) is used to detect such cases with the following parameters:

Match=2, Mismatch=7, Delta=7, PM=80, PI=10, Minscore=50, MaxPeriod=500

#### (5) Read mapping

The existence of a cycle in a de Bruijn graph is a necessary but not a sufficient condition for the existence of tandem repeats in the genome. That is, even if a cycle exists in the de Bruijn graph, the tandem repeat indicated by the cycle may not exist. This is due to the fact that sequences are fragmented. The smaller the value of *k,* the more likely false positive sequences are to appear. This is the disadvantage of using smaller values of *k.* Therefore, read mapping is performed to confirm whether the detected repeat sequences are present in the short reads. In addition to the fact that short read datasets usually have mean coverage depths ≥1, reads that are mapped to repeat sequences are overrepresented in the sequence data compared to other unique sequences. Therefore, to reduce the computational complexity, down-sampling is performed from short reads to align reads so that coverage is 0.1×. It is known that the relationship of *k,* the peak value of *k*-mer occurrences, *P,* and read length can be expressed as Equation 2 (Zerbino and Birney, 2008).

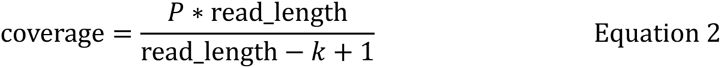

For tandem repeats, alignment is performed for tandemly duplicated unit sequences. Next, sequences that contain unmapped regions are excluded (Figure 5). For alignment, use the “-task megablast” option of blastn.

**Figure 5.**
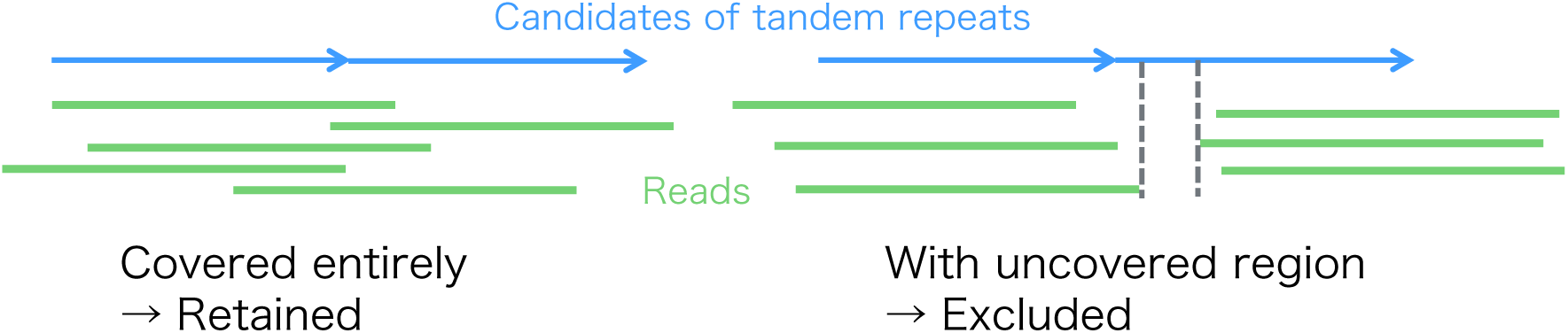
Mapping of reads to tandem repeats.

#### (6) Copy number estimation of detected sequences

For sequences detected by cycle search and after filtering, the copy number is estimated from the configured *k*-mer. A tandem repeat unit of length *u* contains *u k*-mers, and a mean occurrence value is used as a tentative copy number of the sequence (Figure 6). This is used to determine the representative sequence for clustering.

**Figure 6.**
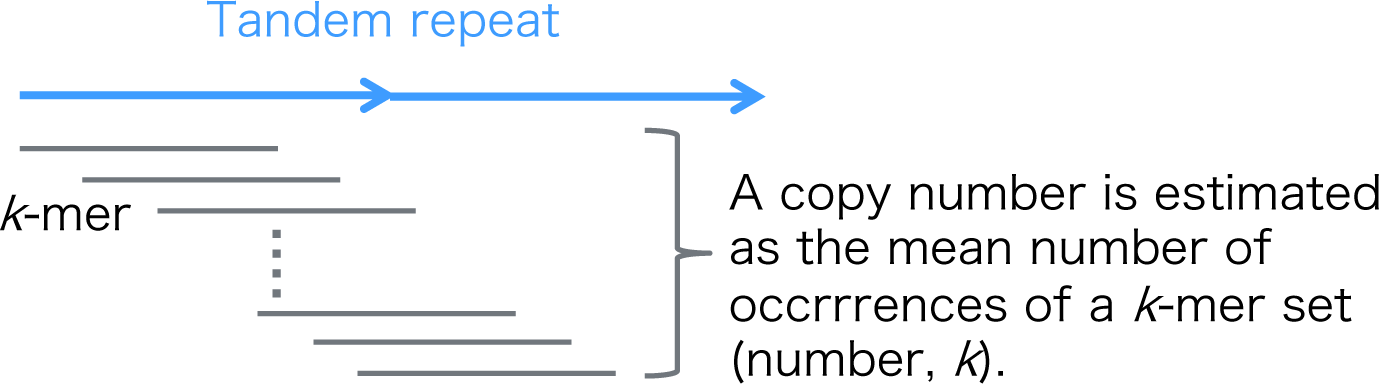
Copy number estimation for tandem repeats.

### Clustering of tandem repeats and copy number estimation

Clustering is performed in two steps: tandem repeats detected from de Bruijn graphs are output as different sequences if they contain even a single nucleotide difference, so that a large number of sequences are detected from sequences in the same cluster. Clustering of sequences with few such differences is performed using CD-HIT, an existing clustering tool. Then, copy number estimation is performed on the obtained representative sequences, and further clustering is performed using further clustering by taking alignment between representative sequences with BLAST. A schematic diagram of the clustering is shown in Figure 7.

**Figure 7.**
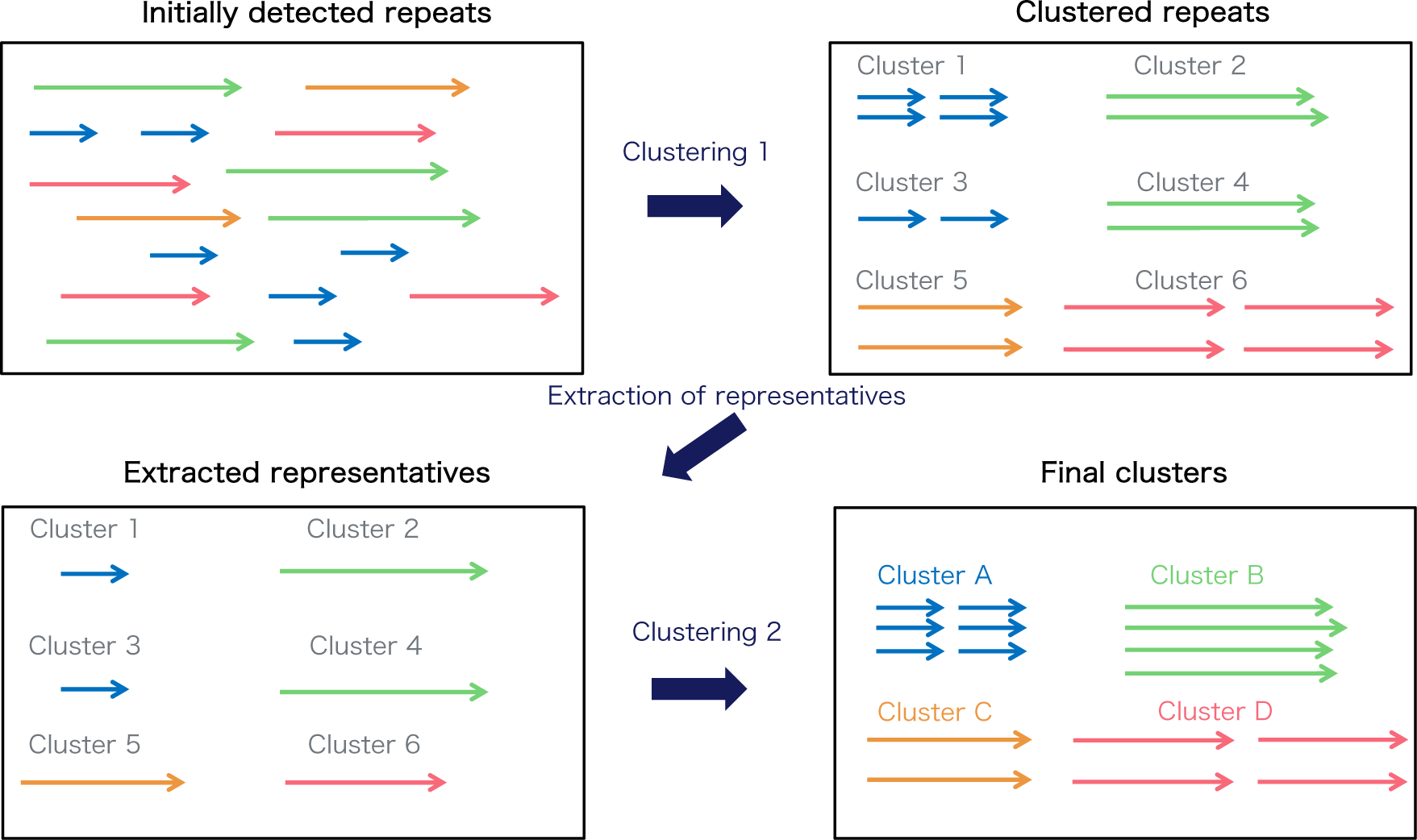
Two-step clustering.

Clustering 1 is done by clustering using CD-HIT and clustering 2 is done by clustering using BLAST.

#### (1) Clustering using CD-HIT

In tandem repeats, there are a number of cases in which the length of one unit depends on the unit length, depending on where the first nucleotide is taken in the tandem repeat. In addition, some similar sequences have different lengths, and in such cases, the sequences may not be clustered if the sequences are connected in tandem and then CD-HIT is used. Therefore, the sequence with the largest copy number among those clustered in tandem is used as the representative sequence, and CD-HIT is used again after restoring the tandem sequences to their original lengths. The following options are used.

word length=3, sequence identity=10, alignment coverage=0.7 (for tandem), 0.8 (for non-tandem), accurate mode, use global identity

#### (2) Estimated number of copies of tandem repeats

Copy number estimation for each cluster. Although the copy number of each sequence can be estimated with the copy number (tentative) obtained in the copy number estimation of the detected sequences, the copy number of the entire cluster cannot be obtained. This is because taking the sum of the copy numbers of the sequences in the cluster will result in double counting of *k*-mer because many of the same *k*-mer are included. Therefore, the copy number in a cluster is estimated by aligning the *k*-mer to the representative sequence.

The representative sequences of each cluster are linked in tandem, and the extracted high-frequency *k*-mer sequences are used as a database for alignment. The copy number is estimated by dividing the sum of the number of occurrences of *k*-mer aligned to the representative sequence by the unit length (Figure 6). This approach is based on the idea of estimating genome size from the distribution of *k*-mer frequencies. When the same *k*-mer is aligned to multiple sequences, the number of occurrences of the *k*-mer is divided by the ratio of the tentative copy number of the aligned sequences. *k*-mer alignment is performed because not all tandem repeats in the genome are detected in the cycle search, The blast option is -task blastn-short-evalue 500-dust no. From the alignment obtained, the alignment is performed at least 0.75 times the length of the *k*-mer, *k.* Since e-value is the expected value of alignment by chance, the larger the number of databases, the larger the value of e-value is likely to be. Therefore, the e-value is set to a large value because the number of *k*-mer databases is particularly large.

#### (3) Clustering using BLASTN

The tandem repeats detected include Figure 8. The tandem repeats detected include sequences that are connected in tandem by a sequence of homologous sequences, as shown in Figure 8, and the question is whether to consider this sequence as a single unit based on its overall length or as a unit by dividing it. In contrast, this method adopts the one with the highest copy number (tentative). For this purpose, those with widely different lengths must also be clustered. However, CD-HIT only clusters sequences up to 70% of the length of the cluster representative sequence. Therefore, in this method, we implemented a further clustering method by aligning representative sequences using blast.

**Figure 8.**
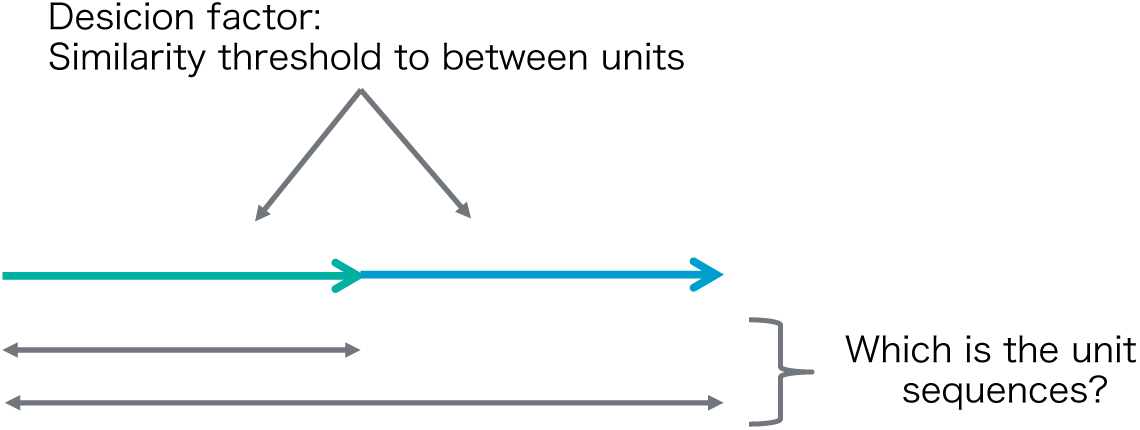
Identification of tandem repeats.

For the above reasons, clustering is performed using not only CD-HIT but also BLAST. The clusters are aligned, and the clusters that are well aligned are combined into the same cluster, and the copy number is calculated as the sum of the copy numbers of the clusters. The option for blastn is-task blastn-short-dust no.

### Detection of interspersed repeat sequences

Since the *k*-mer not used in the detection of tandem repeats is considered to be a *k*-mer derived from interspersed repeats, these *k*-mer are linked together to detect interspersed repeats.

Among the high-frequency *k*-mers, *k*-mers that are considered to be derived from sporadic repeats are obtained, excluding *k*-mer that are aligned with tandem repeats in the copy number estimation of tandem repeats. Using these *k*-mers, we consider the de Bruijn graph and search for pathways constituting the sporadic repeats. The path search method is a depth-first search, and as in the cycle search, the following parameters are set for the search.

*L*: Upper limit of interspersed repeat length to be detected (default value: 5,000)

*N*: Upper limit on the number of nodes to search (default value: 10,000)

*D*: Upper limit of branching depth to search (default value: 5)

The following is a description of the routing procedure (Figure 9).

i. A *k*-mer with zero incoming edges and one or more outgoing edges is extracted and designated as the terminal *k*-mer.
ii. Depth-first search is performed starting from the terminal *k*-mer. The order of search in branching is such that the node with the highest number of occurrences of the next *k*-mer is given priority in the search. When a node with zero outgoing edges is reached, it is stored as a interspersed repeat.
iii. If all paths are explored or if any of *L, N,* or *D* are crossed, the search is terminated and the starting node is removed from the graph.
iv. The search is performed with the next *k*-mer as the starting point.

**Figure 9.**
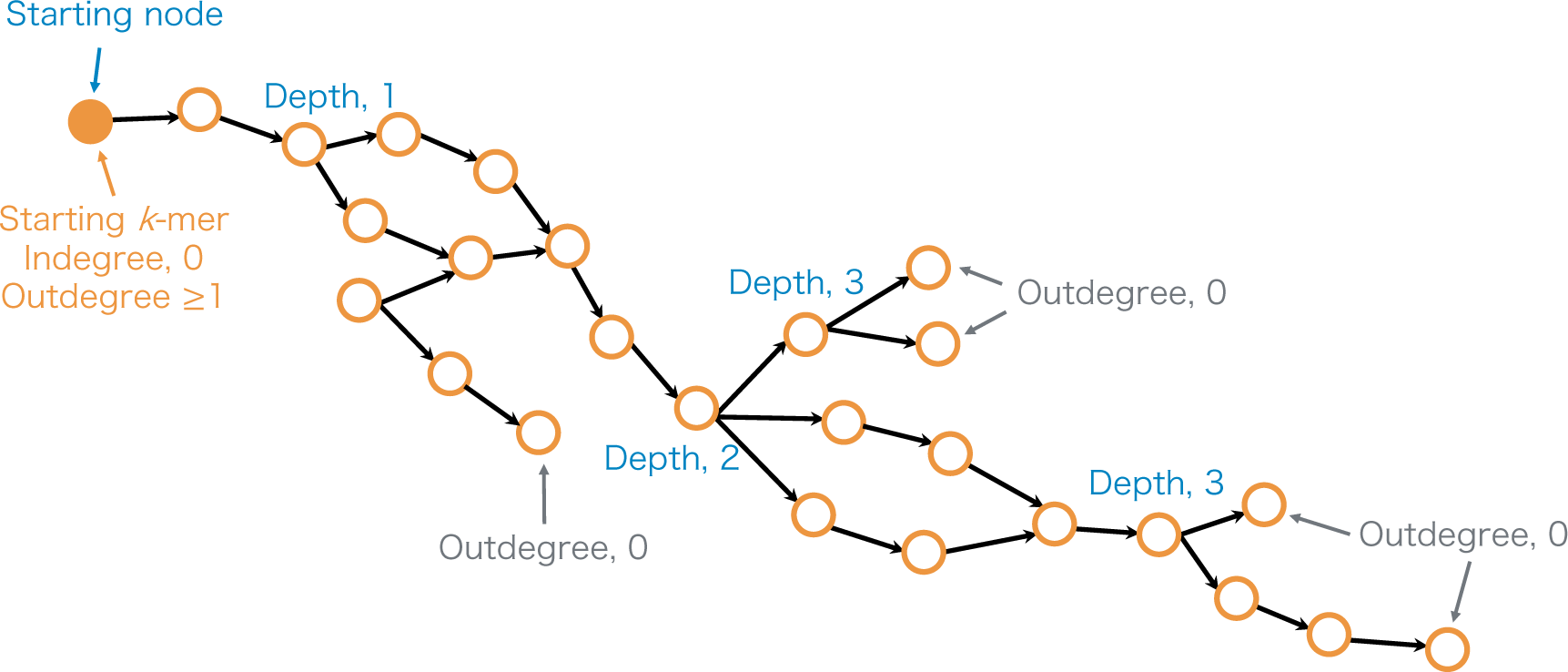
Path finding in interspersed repeats.

#### (1) Read mapping

Reads are mapped to detected interspersed repeat sequences. Since the detected sequences may be longer than the consensus sequence, and the reads used are downsampled, edge region (10% of the sequence length) should not have to be mapped (Figure 10).

**Figure 10.**
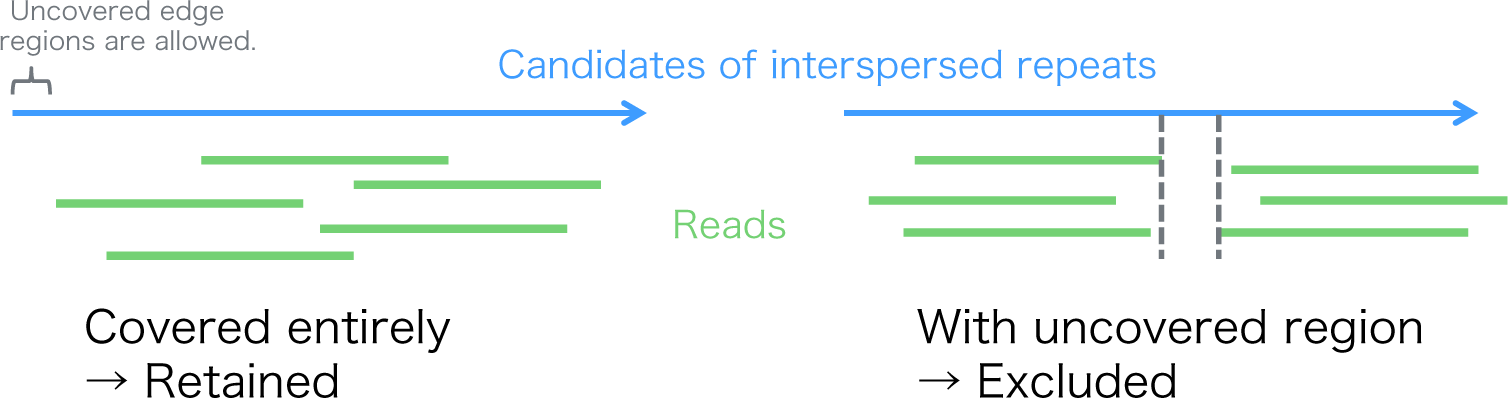
Mapping of reads to interspersed repeats.

#### (2) Copy number estimation of detected sequences

The average number of occurrences of (*u*−*k*+1) different *k*-mer from the detected sequence length *u* is used as a provisional estimate of the copy number (Figure 11).

**Figure 11.**
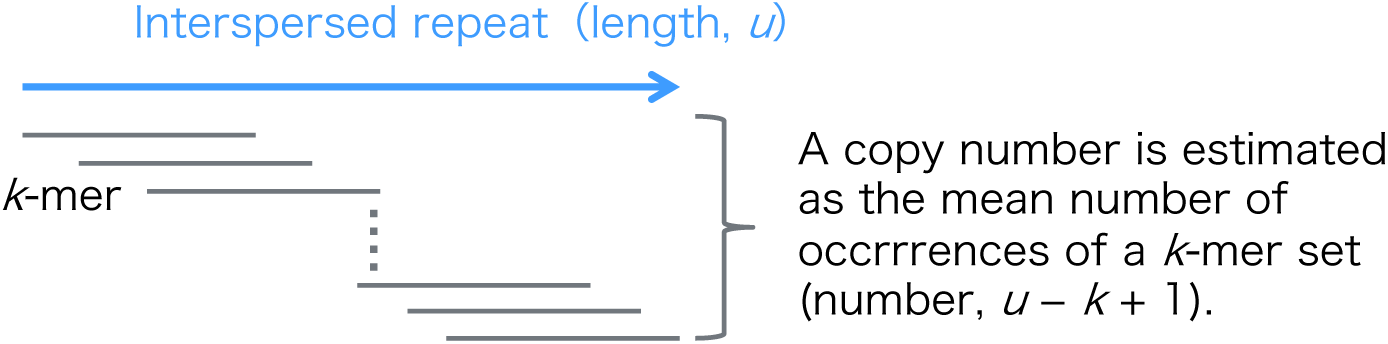
Copy number estimation for interspersed repeats.

#### (3) Clustering of interspersed repeats and copy number estimation

The same procedure as for tandem repeats is used for two-step clustering and copy number estimation (Figure 7).

#### (4) Clustering using CD-HIT

Clustering is performed using CD-HIT and the sequence with the largest estimated copy number is the representative sequence. The following parameters are used:

word length=3, sequence identity=10, alignment coverage=0.8, accurate mode, using global identity

#### (5) Copy number estimation for interspersed repeats

Align the *k*-mer to the representative sequence as in tandem repeat copy number estimation, and estimate the copy number from the *k*-mer to be aligned.

#### (5) Clustering using BLASTN

The obtained clusters are aligned with each other, the aligned clusters are grouped together as the same family, and the sequence with the largest estimated copy number is designated as the representative sequence.

## Results

In the repeat detection tool we developed this time, we confirmed the accuracy of four methods. The repeat sequences detected by this method target regions that are difficult to assemble and do not appear in reference genomes. Therefore, it is difficult to evaluate the accuracy of detection and copy number because there is no set of correct answers. Therefore, we evaluated the method from various aspects using simulated and real data.

### Simulation using E. coli reference genome

In the compare mode of this method, the accuracy of tandem repeat and interspersed repeat detection and copy number estimation are evaluated using simulated data after creating a set of correct answers.

The reference genome of *Escherichia coli* SE11 downloaded from NCBI was used (Oshima *et al*., 2008) (Table 1). First, repeat sequences that had no homology to the reference sequence in SE11 were randomly generated. The lengths were {13 bp, 43 bp, 73 bp, 253 bp, 493 bp, 653 bp, 973 bp}. Mutations were randomly inserted into the original repeat sequence and doubled until mutations of *r* = {0%, 2%, 4%, 6%, 8%, 10%} of the unit length were inserted (Figure 12). Repeats were inserted into the SE11 reference genome and short reads were generated using the ART simulator (Huang *et al*., 2012) ART options were set to a paired-end read length of 100 bp, standard deviation of read length of 10 bp, insert size of 300 bp, read depth of 40x, and HiSeq 2500. We verified that the inserted repeat sequence could be detected by comparing reads before and after repeat insertion (Figure 13). For tandem repeats, 1,000 copies were connected in tandem and inserted at random positions, and for sporadic repeats, 1,000 copies of each sequence were inserted at random positions.

**Figure 12.**
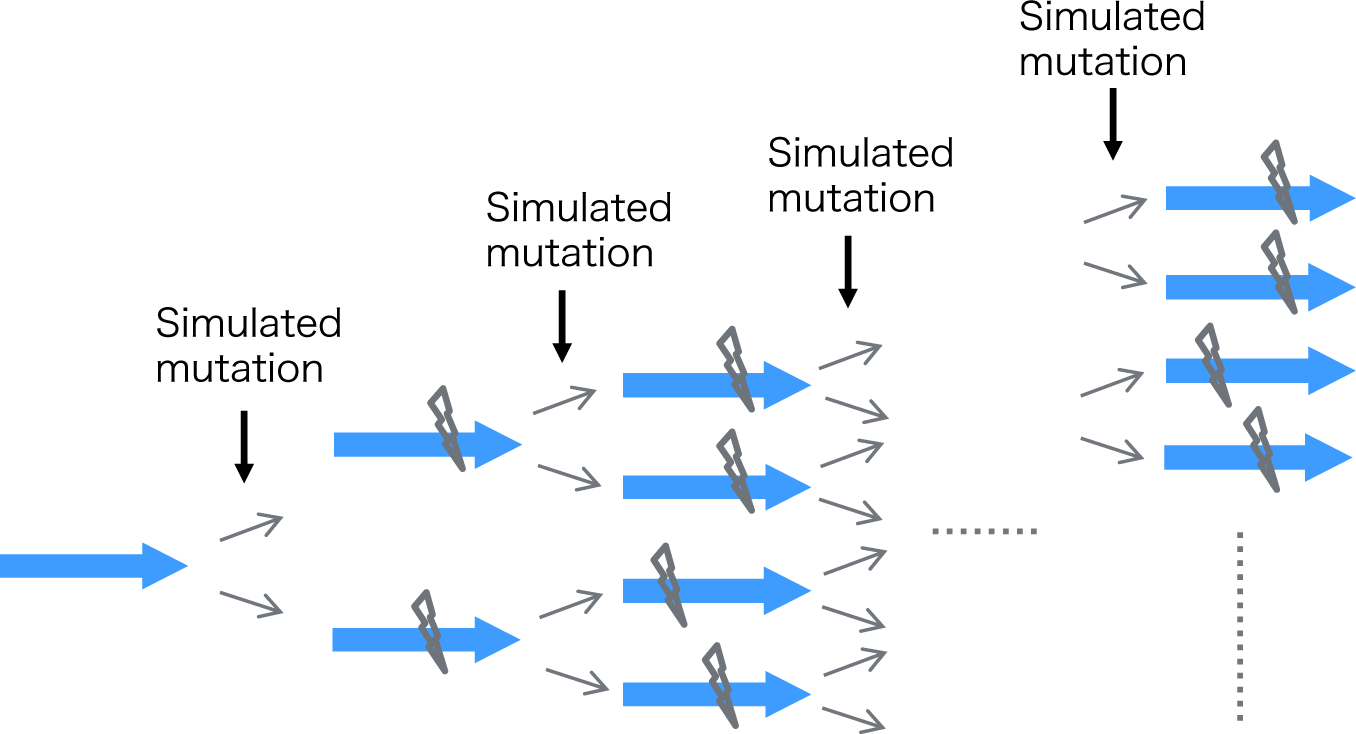
Procedure for creating a repeat sequence with mutations.

**Figure 13.**
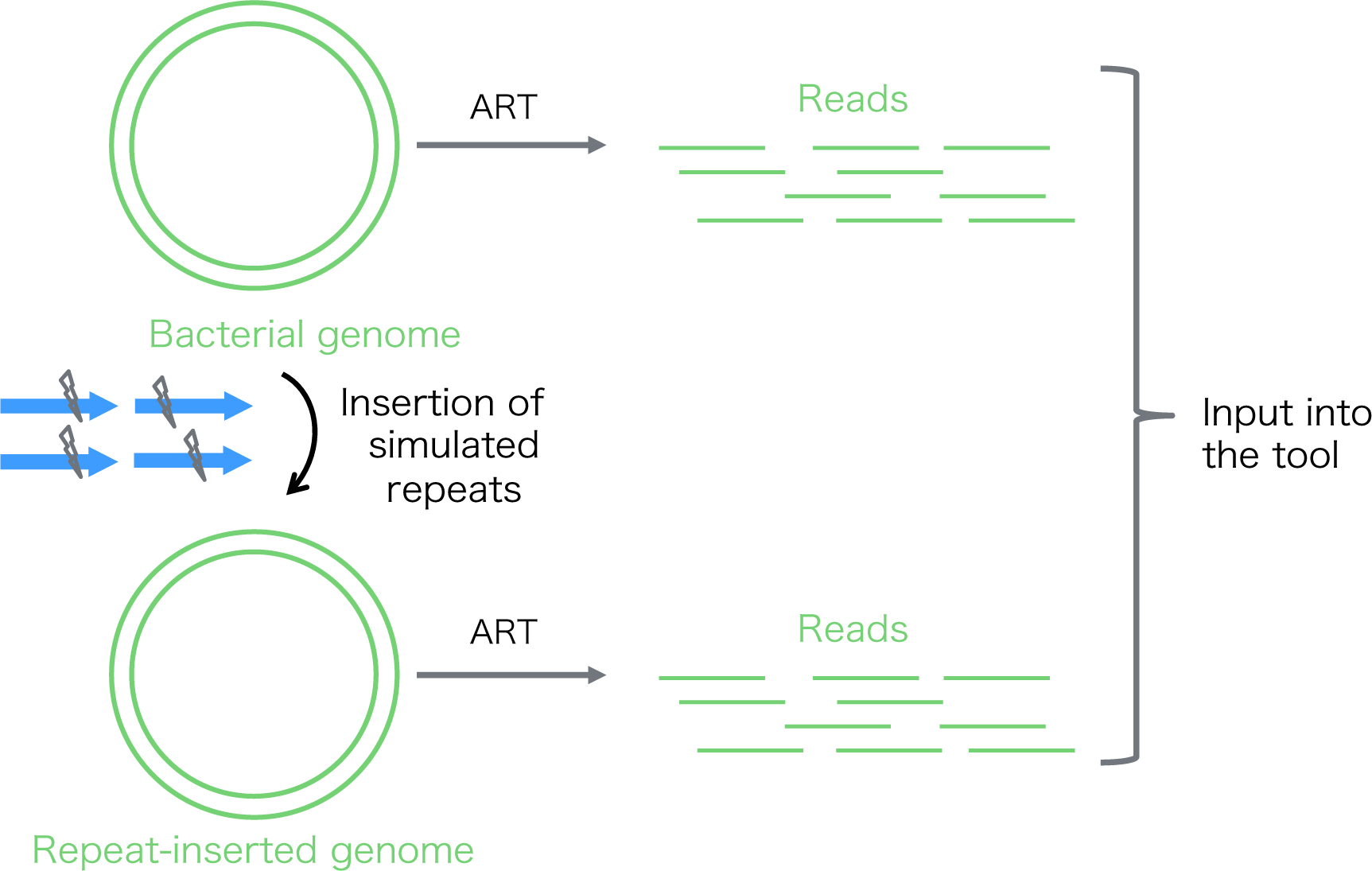
Schematic model of benchmarking with *E. coli* simulation data. ART: read simulator.

**Table 1.**
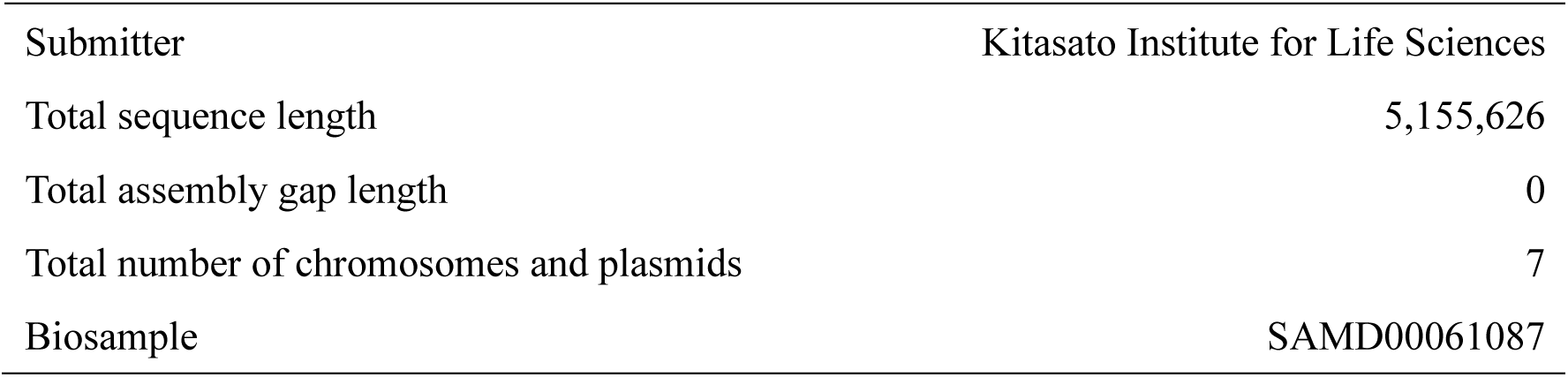
Reference genome information for *E. coli* SE11.

|  |  |
| --- | --- |
| Submitter | Kitasato Institute for Life Sciences |
| Total sequence length | 5,155,626 |
| Total assembly gap length | 0 |
| Total number of chromosomes and plasmids | 7 |
| Biosample | SAMD00061087 |

The results for tandem repeats and interspersed repeats are respectively Figure 14 and Figure 15 respectively. In the tandem repeat benchmark, the inserted sequences were successfully detected except in the case of 10% mutation rate and unit length, 73 bp, and in all these cases the estimated copy number was in the range of 500–1100 bp. The inserted sequences are successfully detected except in the cases of 8% and 10% mutation rates and 43 bp unit length for the interspersed repeat benchmarks, and the estimated copy number is within the range of 500–1100 bp in all cases where the mutation rate is 6% or less. Thus, it is suggested that when each repeat sequence differs from the consensus sequence by 6% or less, cycle_finder can successfully detect the unit sequence and estimate the copy number with an error within 50% of the true value. It was also suggested that when the unit sequence length is longer than 253 bp, cycle_finder works well even if the mutation rate is as large as 10%.

**Figure 14.**
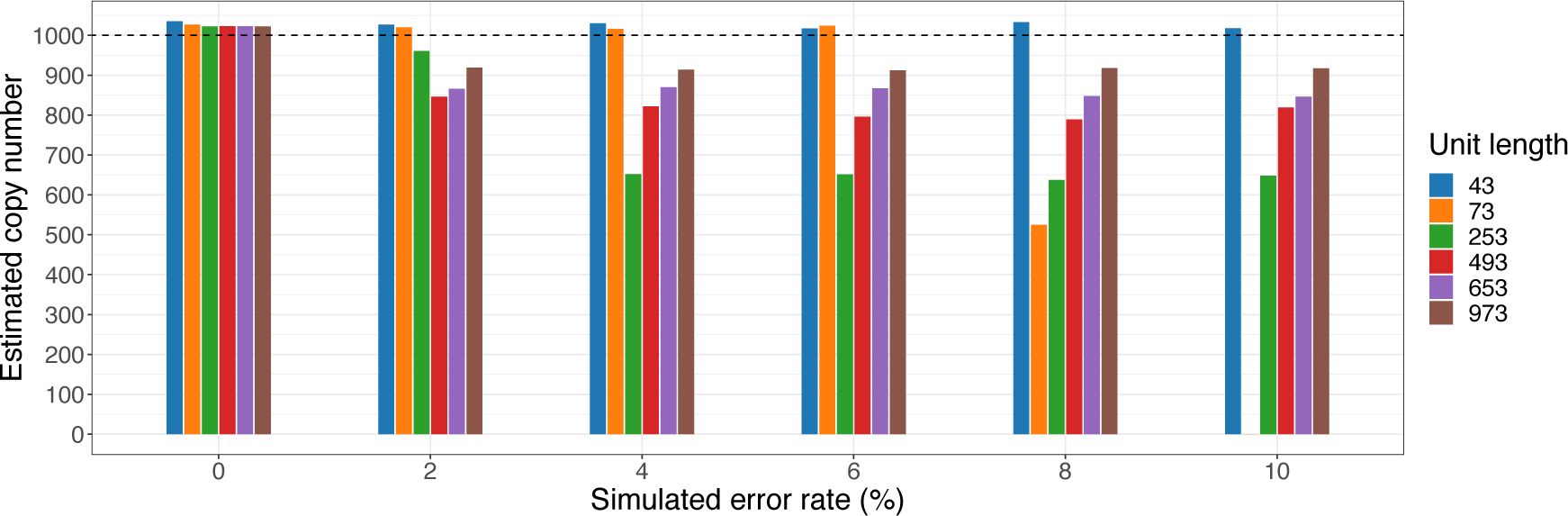
Benchmarks of copy number estimation for tandem repeats. Horizontal axis, number of mutations relative to unit length; vertical axis, estimated number of copies.

**Figure 15.**
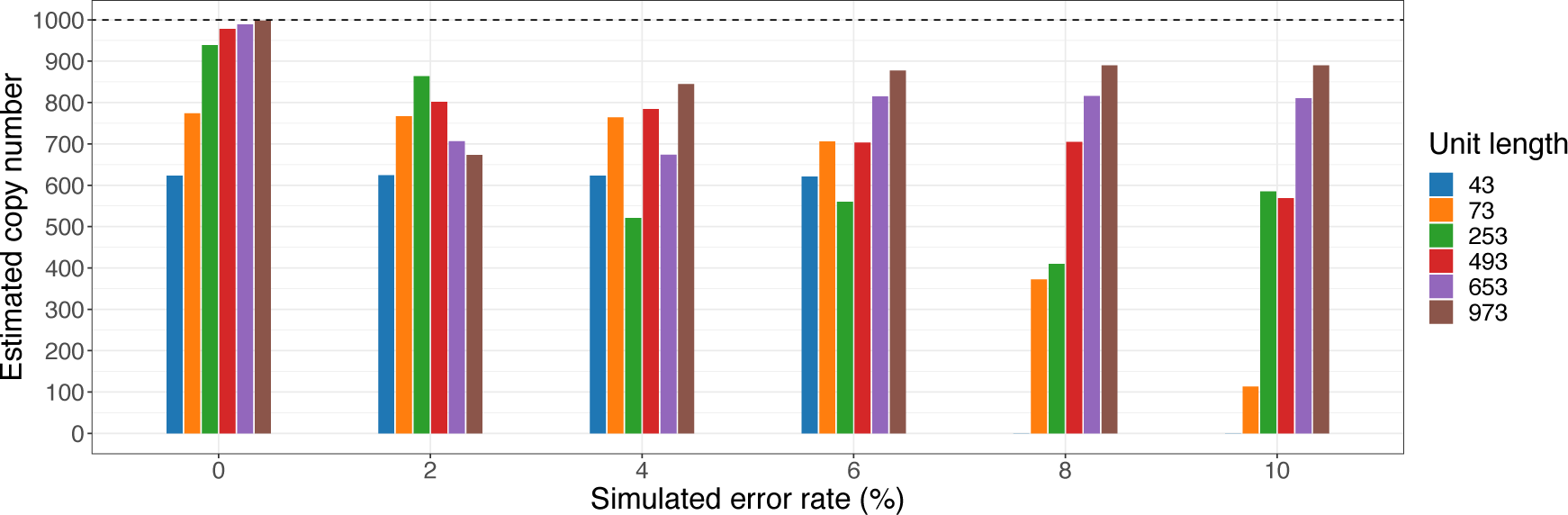
Benchmarks of copy number estimation for interspersed repeats. Horizontal axis, number of mutations relative to unit length; vertical axis, estimated number of copies.

### Benchmark using human CHM13 data

The human (*Homo sapiens*) hydatidiform mole cell, CHM13, can be considered a pseudo-monoploid state, and a complete reference genome sequence with no gaps has been published (Nurk *et al*., 2022) Therefore, benchmarking can be performed with the reference genome sequence as the correct data. Here, we use Illumina paired-end reads (publicly available data, see Table 2) were input into each tool, the output repeat sequence libraries (representative sequence sets for each repeat sequence class) were aligned to the reference genome sequence, and performance was evaluated based on the overlap between the alignment region and the reference repeat sequence annotation.

**Table 2.**
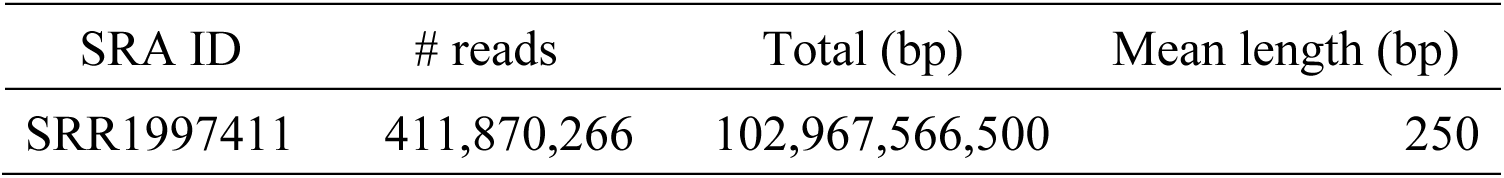
Statistics for human CHM13 reads.

| SRA ID | # reads | Total (bp) | Mean length (bp) |
| --- | --- | --- | --- |
| SRR1997411 | 411,870,266 | 102,967,566,500 | 250 |

The reference genome sequence was T2T-CHM13 v2.0 excluding the Y chromosome, which was from another individual. The total length is 3,054,815,472 bp. The repeat sequence annotation was the RepeatMasker-based one (rm.out) in the RefSeq database. The breakdown of the classifications of the elements of the repeat annotations is shown in Table 3 shows the breakdown of the classifications of the elements of the repeat annotations.

**Table 3.**
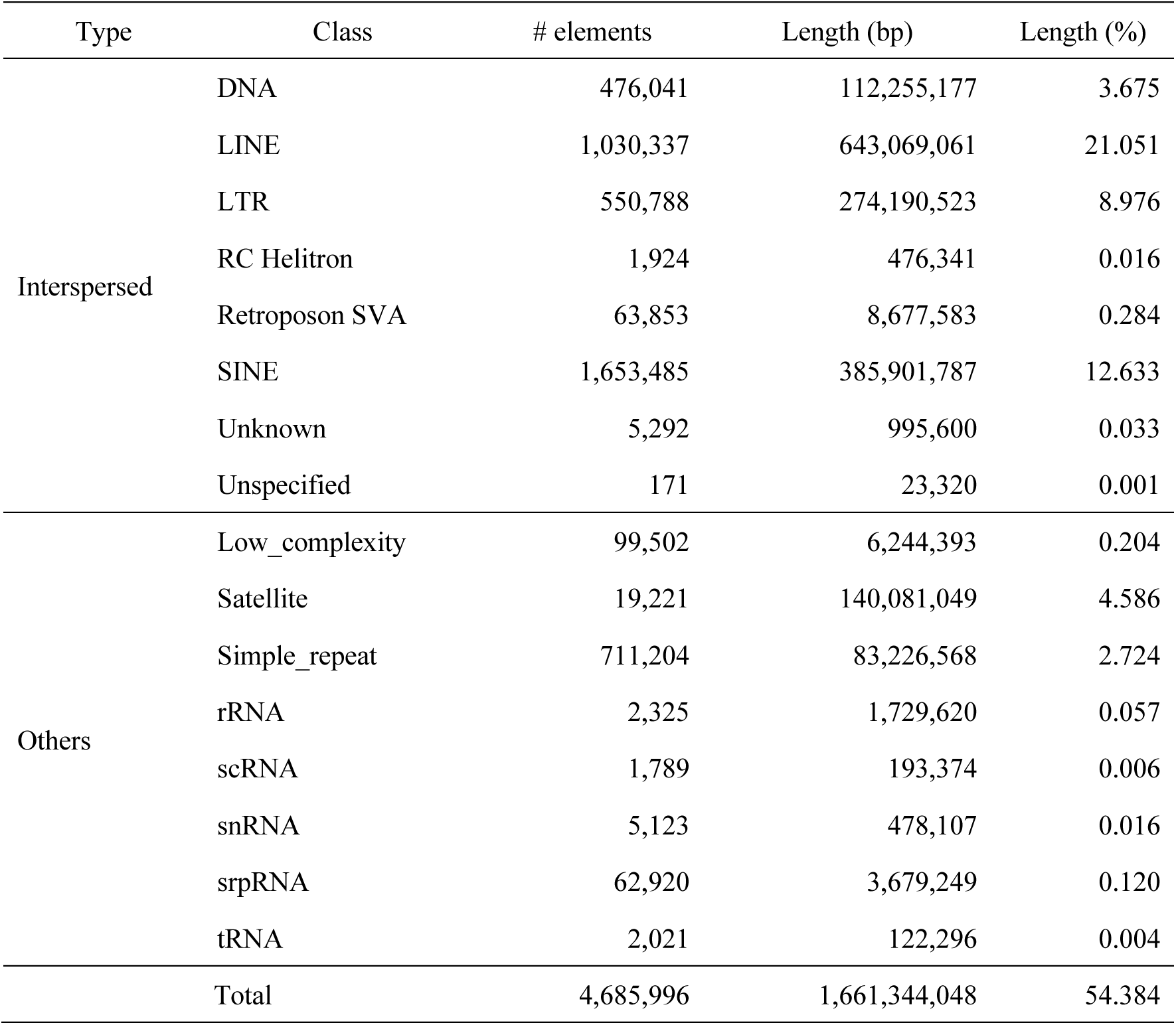
Breakdown of human CHM13 reference repeat annotations.

Benchmarking tools include:

(1) cycle_finder (v1.0.0)
(2) TAREAN (v0.3.8.1)
(3) MEGAHIT (v1.1.3) + RepeatModeler (v2.0)
(4) Platanus (v1.2.4) + RepeatModeler (v2.0)

Since TAREAN recommends that reads be run downsampled so that the average coverage depth <1, we tested four different settings for the amount of input reads: no downsample (full), average coverage depth 0.5×, average coverage detph 0.1×, and number of reads 500k (-s 500k). For (3) and (4), we used a general *de novo* assembler: MEGAHIT (Li *et al*., 2015), Platanus (Kajitani *et al*., 2014) and then run the *de novo* repeat library construction tool, RepeatModeler (Flynn *et al*., 2020) on the resultant sequence. These are to be compared with (1) and (2), which are specialized to repeat sequences.

stats of the output array set foreach tool. Table 4 shows the statistics of the output sequence set for each tool. Note that the phiX sequences have been removed (aligned to the phiX reference sequence in BLASTN and removed if identity ≥95 and query-coverage ≥50% and alignment length ≥17).

**Table 4.** Statistics of output sequence set (Human CHM13).

| Tool | #<br>sequences | Total (bp) | Minimum<br>length (bp) | Mean<br>length (bp) | Maximum<br>length (bp) |
| --- | --- | --- | --- | --- | --- |
| cycle_finder | 3,555 | 153,524 | 1 | 43 | 9,522 |
| cycle_finder (tandem only) | 97 | 6,223 | 1 | 64 | 555 |
| TAREAN (full) | 17 | 30,090 | 15 | 1,770 | 16,569 |
| TAREAN (0.5×) | 17 | 30,038 | 15 | 1,767 | 16,569 |
| TAREAN (0.1×) | 13 | 23,643 | 15 | 1,819 | 16,569 |
| TAREAN (-s 500k) | 10 | 2,012 | 15 | 201 | 994 |
| MEGAHIT + RepeatModeler | 726 | 500,023 | 34 | 689 | 5,910 |
| Platanus + RepeatModeler | 1,244 | 496,613 | 29 | 399 | 4,680 |

For cycle_finder, all output sequence sets and those output as tandems are noted separately. TAREAN is characterized by a small number of output sequences in any setting. megahit, Platanus + RepeatModeler’s total is large but accurate. The evaluation of the accuracy is discussed below.

Using the reference repeat annotation as the ground-truth data, the following procedure was used to calculate the region length and based recall and precision.

(1) Align the output sequence set of each tool to the reference genome sequence (tool, BLASTN; options, “ -task blastn -dust no -qcov_hsp_perc 50 -perc_identity 80 -evalue 0.1 -word_size 5 - max_target_seqs 10000000”). Sequences output as tandem repeats are duplicated until they reach 20 bp or more.
(2) Calculate recall and precision for alignment regions. The correct solution and the region to be evaluated are merged without overlap using bedtools merge. Recall is calculated as the overlap region length between the alignments and the ground-truth regions divided by the ground-truth region length. Precision is calculated as the overlap region length between the alignments and the ground-truth regions divided by the alignment region length. We also performed benchmarking using repeat elements with divergences ≤20% as the ground-truth data. This is intended to exclude elements that contain a lot of mutations and are not substantially repeat sequences. The results are shown in Table 5.

**Table 5.** Human CHM13 benchmark result based on alignment length.

| Ground-truth set | Tool | Recall. | Precision | F1 |
| --- | --- | --- | --- | --- |
| all | cycle_finder | 0.431 | 0.988 | 0.600 |
|  | cycle_finder (tandem only) | 0.163 | 0.994 | 0.280 |
|  | TAREAN (full) | 0.085 | 0.990 | 0.157 |
| | TAREAN (0.5×) | 0.090 | 0.990 | 0.164 |
| | TAREAN (0.1×) | 0.097 | 0.994 | 0.177 |
|  | TAREAN (-s 500k) | 0.080 | 0.993 | 0.148 |
|  | MEGAHIT + RepeatModeler | 0.263 | 0.987 | 0.415 |
|  | Platanus + RepeatModeler | 0.293 | 0.982 | 0.452 |
| divergence $\leq 20$ | cycle_finder | 0.652 | 0.934 | 0.768 |
|  | cycle_finder (tandem only) | 0.255 | 0.971 | 0.404 |
|  | TAREAN (full) | 0.135 | 0.983 | 0.238 |
| | TAREAN (0.5×) | 0.142 | 0.984 | 0.249 |
| | TAREAN (0.1×) | 0.156 | 0.991 | 0.269 |
|  | TAREAN (-s 500k) | 0.128 | 0.990 | 0.226 |
|  | MEGAHIT + RepeatModeler | 0.417 | 0.976 | 0.584 |
|  | Platanus + RepeatModeler | 0.458 | 0.957 | 0.620 |

cycle_finder achieves the largest recall, F1, in both conditions, with a precision of 0.93 or higher. TAREAN has a smaller recall, and the RepeatModeler-based results with larger totals also show that recall is inferior to cycle_finder.

However, the alignment length-based precision has a disadvantage as an evaluation index, since it is not reflected in the evaluation even if sequences that do not align to the reference genome are output, and even if non-repeat sequences are output, they are not reflected because they are small as the alignment region length. Therefore, we calculated the precision based on the number of sequences, using the sequences that have multi-hit to the reference genome sequence as the output sequences of each tool (Table 6). The alignment method and hit conditions are the same as those described above (tool, BLASTN), and sequences output as interspersed repeats that are less than 20 bp were excluded from the evaluation. This is because the e-value >0.1 for an exact match.

**Table 6.**
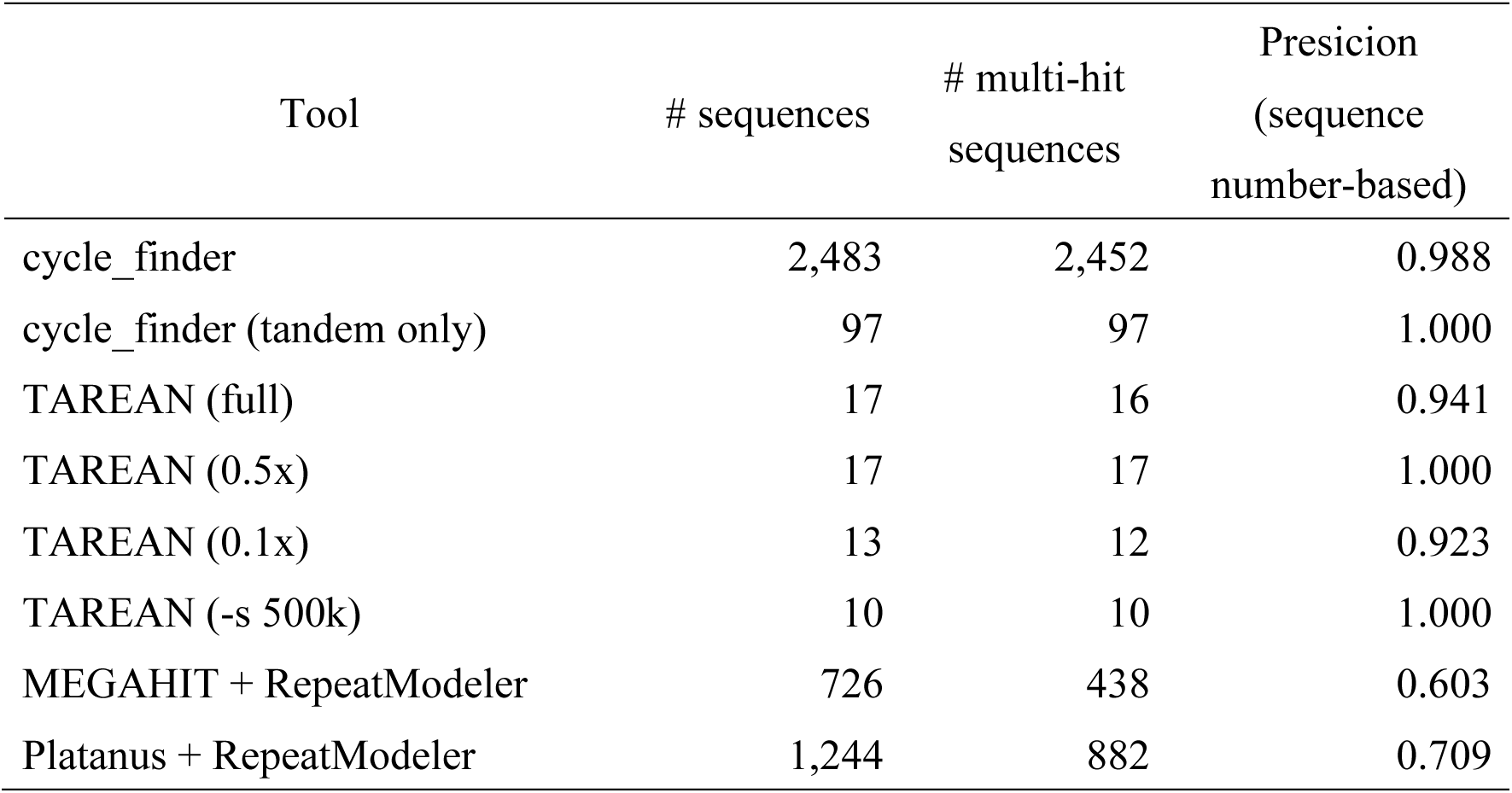
Human CHM13 sequence number-based precision.

| Tool | # sequences | # multi-hit<br>sequences | Precision<br>(sequence<br>number-based) |
| --- | --- | --- | --- |
| cycle_finder | 2,483 | 2,452 | 0.988 |
| cycle_finder (tandem only) | 97 | 97 | 1.000 |
| TAREAN (full) | 17 | 16 | 0.941 |
| TAREAN (0.5x) | 17 | 17 | 1.000 |
| TAREAN (0.1x) | 13 | 12 | 0.923 |
| TAREAN (-s 500k) | 10 | 10 | 1.000 |
| MEGAHIT + RepeatModeler | 726 | 438 | 0.603 |
| Platanus + RepeatModeler | 1,244 | 882 | 0.709 |

The results show that the precision of cycle_finder, TAREAN is large (>0.98), while the RepeatModeler-based output is small (<0.71). Although the total sequence set was large, it is suggested that a large number of non-repeating sequences were also output.

As an overall evaluation, plots for recall (based on region length) and precision (based on number of sequences) are shown in Figure 16.

**Figure 16.**
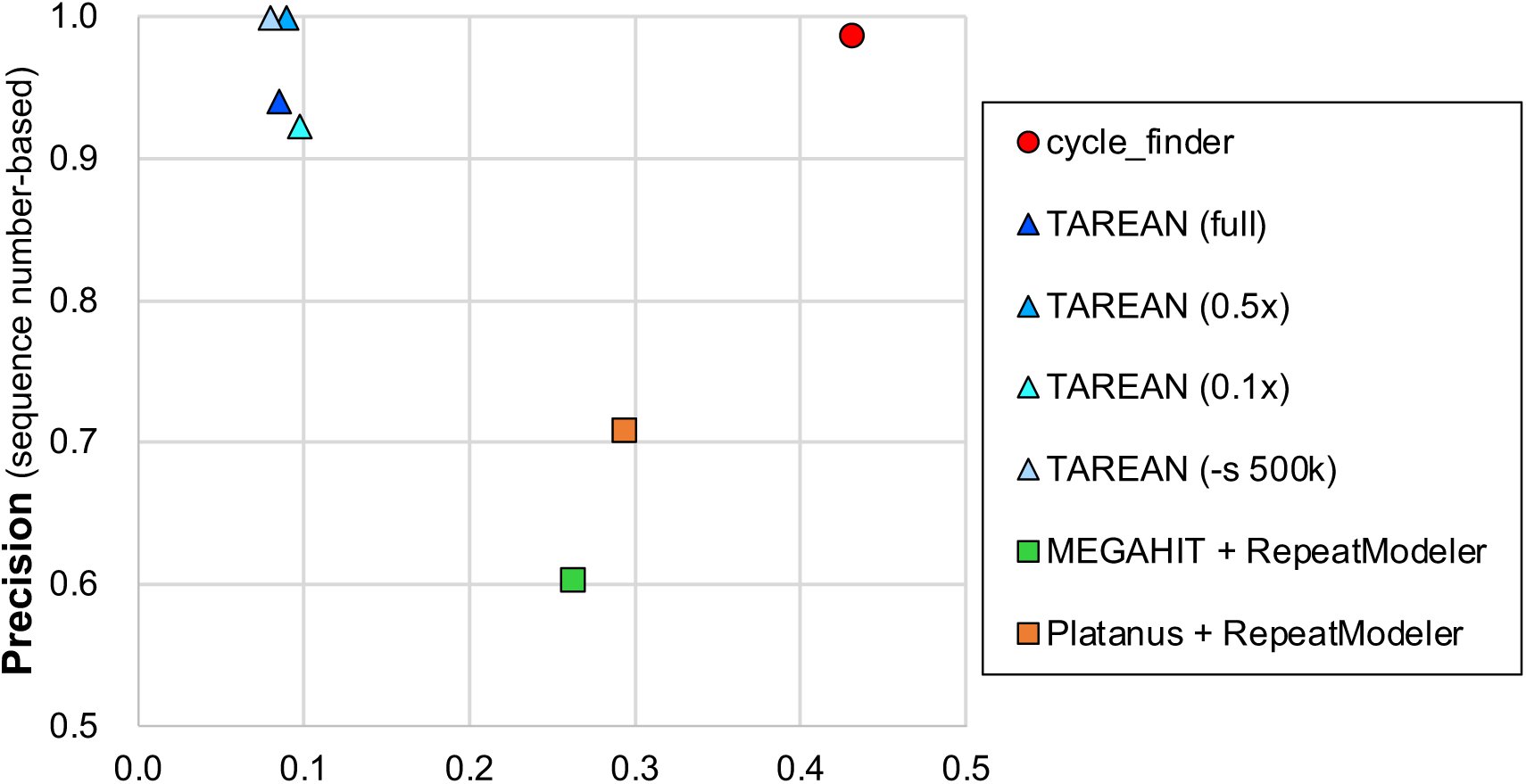
CHM13 recall-precision plot.

A point located in the upper right corner indicates higher overall performance, while cycle_finder is shown to have high performance with both values being large.

The following procedure was used to investigate the classification of cycle_finder output sequences:

(1) Representative sequences from the tool output were aligned to the reference genome.
(2) Top hits (bit-score criteria) of representative sequences were determined.
(3) If there was overlap with the reference annotation, it was mapped. If there were multiple candidates due to multiple hits, the top of the total overlap length was adopted.

The classification breakdown of the output tandem and interspersed sequences are shown in Table 7 and Table 8, respectively.

**Table 7.** Classification breakdown of tandem repeat output from cycle_finder (human CHM13).

| Repeat class | count | rate (%) |
| --- | --- | --- |
| Simple_repeat | 46 | 47.42 |
| Satellite | 9 | 9.28 |
| no_hit | 9 | 9.28 |
| SINE/Alu | 6 | 6.19 |
| LTR/ERV1 | 6 | 6.19 |
| LINE/L1 | 6 | 6.19 |
| Satellite/centr | 5 | 5.15 |
| Retroposon/SVA | 3 | 3.09 |
| transfer RNA | 1 | 1.03 |
| Satellite/acro | 1 | 1.03 |
| rRNA | 1 | 1.03 |
| LTR/ERVL-MaLR | 1 | 1.03 |
| LTR/ERVK | 1 | 1.03 |
| Low_complexity | 1 | 1.03 |
| DNA/hAT-Charlie | 1 | 1.03 |

**Table 8.** Classification breakdown of interspersed repeat output from cycle_finder (human CHM13).

| Repeat class | count | rate (%) |
| --- | --- | --- |
| LINE/L1 | 1,166 | 33.72 |
| no_hit | 1,110 | 32.10 |
| SINE/Alu | 342 | 9.89 |
| LTR/ERVL-MaLR | 271 | 7.84 |
| LTR/ERV1 | 139 | 4.02 |
| DNA/TcMar-Tigger | 106 | 3.07 |
| Satellite/centr | 72 | 2.08 |
| Simple_repeat | 58 | 1.68 |
| LTR/ERVL | 47 | 1.36 |
| DNA/hAT-Charlie | 46 | 1.33 |
| LTR/ERVK | 23 | 0.67 |
| Retroposon/SVA | 16 | 0.46 |
| LINE/L2 | 14 | 0.40 |
| Satellite | 13 | 0.38 |
| SINE/MIR | 11 | 0.32 |
| Low_complexity | 7 | 0.20 |
| DNA/TcMar-Mariner | 6 | 0.17 |
| srpRNA | 5 | 0.14 |
| rRNA | 2 | 0.06 |
| DNA/PiggyBac | 2 | 0.06 |
| DNA/MULE-MuDR | 1 | 0.03 |
| DNA/hAT-Tip100 | 1 | 0.03 |

Simple_repeat and Satellite were more common in the tandem repeat output, and transposons such as LINE, SINE, and LTR were more common in the interspersed repeat output, suggesting that cycle_finder worked as designed.

### Application to the analysis of somatic chromosome diminution in *Ascaris suum*

Tandem repeats have also been found to be included in deletion sequences in chromosome in the roundworm (*Ascaris suum)* (Wang *et al*., 2012). The tandem repeats are highly likely to be deleted in chromosome diminution in *A. suum*. In particular, previous studies have shown that a satellite sequence (unit length, 121 bp) with a high copy number is deleted in somatic cells. In addition, the copy number of telomeric repeat (unitlength, 6 bp) is also known to differ significantly between germline and somatic cells. Therefore, we applied the compare mode of cycle_finder to the *A. suum* data and verified whether the previously studied sequences could be detected.

We used the compare mode of cycle_finder to detect somatically deleted sequences using paired-end libraries from germline and somatic cells of *A. suum*, which were downloaded from NCBI (Table 9). This data were used in the previous study above (Wang *et al*., 2012).

**Table 9.**
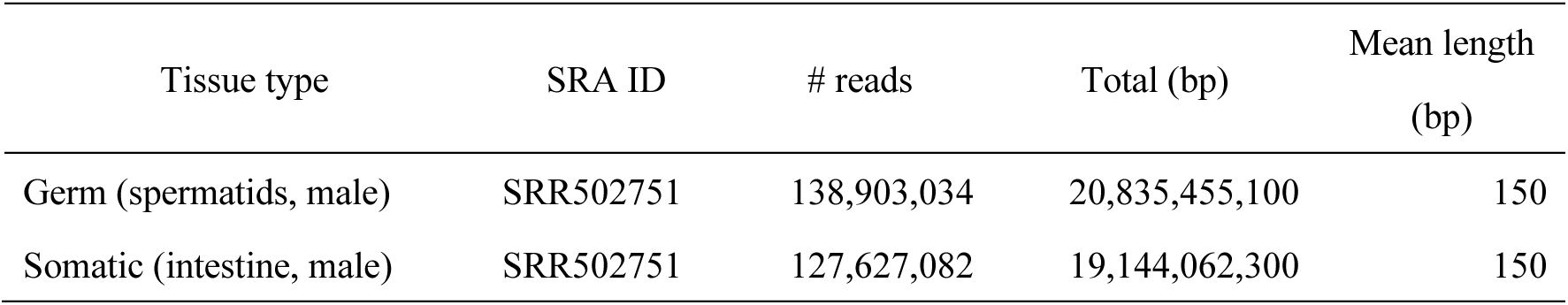
Statistics for *A. suum* reads.

| Tissue type | SRA ID | # reads | Total (bp) | Mean length<br>(bp) |
| --- | --- | --- | --- | --- |
| Germ (spermatids, male) | SRR502751 | 138,903,034 | 20,835,455,100 | 150 |
| Somatic (intestine, male) | SRR502751 | 127,627,082 | 19,144,062,300 | 150 |

The results obtained are shown in Figure 17. A summary of germline and somatic copy numbers is shown in Table 10. We succeeded in detecting the sequences reported in previous studies. The copy number values that differ from those in the previous study are thought to be due to the different copy number estimation methods. The present method uses the frequency of occurrence of *k*-mer to calculate the copy number, whereas the method of the previous study estimates the copy number by mapping reads to repeat sequences. It is difficult to determine which method is correct because it depends on the parameters of alignment and mapping, and it is a question of how much difference in repeat sequences to allow.

**Figure 17.**
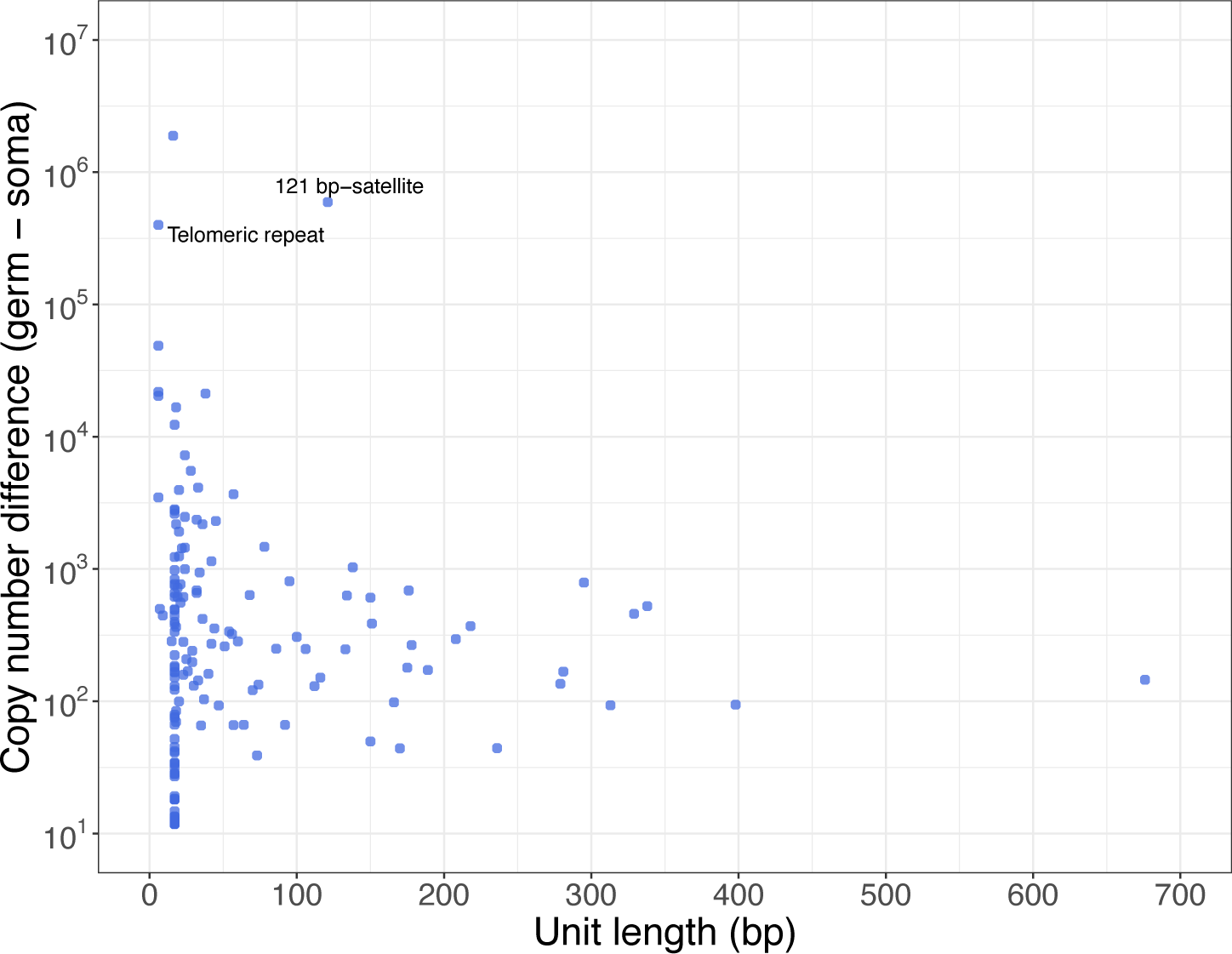
Detection of tandem repeats in *A. suum.* Horizontal axis, repeat length; vertical axis, estimated number of copies (log scale); blue dots, elements detected by cycle_finder.

**Table 10.** Result of cycle_finder for previously reported sequences in *A. suum*.

| Sequence name | Unit length (bp) | Germ copy<br>number | Somatic copy<br>number | Copy number<br>difference |
| --- | --- | --- | --- | --- |
| 121 bp-satellite | 121 | 629,797<br>(245,526) | 35,531<br>(13,478) | 594,266<br>(232,048) |
| Telomeric<br>repeats | 6 | 854,712<br>(297,546) | 455,303<br>(176,990) | 399,409<br>(12,556) |

Values in parentheses are estimates from the previous study.

## Discussion

In this study, we developed a method for constructing repeat unit sequences based on cycle detection in de Bruijn graphs: cycle_finder, using low-cost short read sets as input. This tool not only constructs sequences, but also calculates copy number differences for each sample and between samples based on the number of occurrences of *k*-mer.

Benchmarking with simulated *E. coli* data confirmed the basic performance of the cycle_finder with respect to repeat unit sequence length and mutation rate. It was shown that if the mutation rate is less than 6%, sequence construction is successful for both tandem repeats and interspersed repeats. Furthermore, even if the mutation rate was larger than that, the construction was successful in the case of long unit sequences.

Next, we benchmarked using human CHM13 data to examine its validity on real data. Here benchmarking can be done by using the complete reference data as the correct answer. As a result, cycle_finder showed high values for both recall and precision, while TAREAN, which targets only tandem repeats, resulted in a smaller number of output sequences and smaller recall. The combination of a general-purpose *de novo* assembler and RepeatModeler is designed to output mainly sporadic repeats, and although the total is large, the output contains many non-repeat sequences, resulting in a small precision. Overall, we believe that the real data demonstrates the usefulness of the cycle_finder.

Finally, the usefulness of cycle_finder in the analysis of somatic mutations was tested using data from *A. suum*. By applying this tool, we were able to detect two types of tandem repeats reported in previous studies, which are somatic cell deletions in somatic cells. Several previously unidentified tandem repeats were also detected, suggesting that the comprehensiveness of our tool’s results may be excellent: programmed chromosome reductions have been reported in *A. suum*, and such programmed genome rearrangements have been reported in many lineages. However, the breakdown of the sequences to be deleted has only been examined in a limited number of lineages, such as the roundworms, hagfishes, and Tetrahymena. This tool is expected to be useful as a means of investigating this interesting phenomenon for many non-model organisms.

TAREAN, an existing tool to construct repeat sequence units *de novo*, clusters downsampled short reads based on all-vs-all alignment and detects tandem repeats within each cluster. Since this method takes time for alignment, it is necessary to adjust the downsample ratio, but automatic determination of that value has not been implemented. Therefore, actual application may require multiple trials, which could be time-consuming and labor intensive. cycle_finder bypasses the alignment process by employing a fast de Bruijn graph-based method, allowing readsets to be entered without downsamples. TAREAN also detects cycles in the graph structure, but it does not target the non-cycle parts, i.e., the parts corresponding to interspersed repeats, which may be the reason why recall is smaller than cycle_finder in the benchmark.

Other methods for detecting repeat unit sequences include a combination of a general-purpose *de novo* assembler and a tool for detecting repeat sequences on contigs. It is important to note that many tools do not output the repeat sequences in a summarized format, such as by clustering the sequences. RepeatModeler, which was benchmarked in this study, outputs representative sequences for each repeat class while utilizing a number of tools including TRF. One problem with this strategy is that the full length of the repeat unit sequence may not be constructed during the initial contig construction phase. The diversity of repeat sequences due to their mutations can introduce a number of branching structures into the graph structure (e.g., de Bruijn graph) used in the process. The cycle_finder addresses this problem by assuming the existence of branches in the cycle and performing path finding. In this benchmark on the human genome, the recall of RepeatModeler may have been smaller due to this problem. Another newly discovered problem is that sequences that cannot be considered truly repeats, which do not appear multiple times in the correct reference genome sequence, are output, resulting in a smaller precision. This may be due to the presence of erroneous duplications in the contig or to the loose sequence clustering criteria of RepeatModeler.

One of the challenges of the cycle_finder algorithm is that some parameters are not automatically determined at runtime: the default values of the threshold *α* for the number of copies to extract k-mers, the upper limit for the repeat length, the number of nodes to search, and the upper limit for the depth during path finding were empirically determined based on performance and execution time determined by taking into account the performance and execution time. Automatic determination of these values would improve usability. In addition, although k was set to 17 when constructing the de Bruijn graph, it is expected that in the future performance will be further improved by applying multiple values and integrating the results, as has been demonstrated to be effective in other assemblers.

With respect to repeat sequences, analysis of somatic variation and inter-individual diversity requires comparison of multiple samples. Although long reads are effective for constructing repeat sequences, the cost is greater than short reads. A combined analysis of short reads and cycle_finder is suitable for a low-cost, multiple-sample plan. Another possible procedure would be to conduct the analysis with cycle_finder as a test and attempt to add long-read data when interesting findings are suggested. The application of the tool developed in this study is expected to accelerate research on repeat sequence regions that have been difficult to analyze so far.

## Acknowledgements

We thank Prof. Souichirou Kubota, Dr. Yuji Goto, and Mr. Kohei Nagao for their valuable feedback on the applications of cycle_finder.

## Data availability

cycle_finder is implemented in C++ and is freely available at GitHub (https://github.com/rkajitani/cycle_finder). The reference genomes and sequencing data used in this study are public data, and their accessions and SRA IDs are described in the corresponding sections.

